# Prevalent chromosome fusion in *Vibrio cholerae* O1

**DOI:** 10.1101/2024.06.12.598706

**Authors:** Aline Cuénod, Denise Chac, Ashraful I. Khan, Fahima Chowdhury, Randy W. Hyppa, Susan M. Markiewicz, Stephen B. Calderwood, Edward T. Ryan, Jason B. Harris, Regina C. LaRocque, Taufiqur R. Bhuiyan, Gerald R. Smith, Firdausi Qadri, Patrick Lypaczewski, Ana A. Weil, B. Jesse Shapiro

**Affiliations:** Department of Microbiology and Immunology, McGill University, Montréal, Québec, Canada; Department of Medicine, University of Washington, Seattle, Washington, United States; International Centre for Diarrhoeal Disease Research, Bangladesh, (icddr,b), Dhaka, Bangladesh; Division of Basic Sciences, Fred Hutchinson Cancer Center, Seattle, Washington, United States; Division of Infectious Diseases, Massachusetts General Hospital, Boston, Massachusetts, USA; Department of Medicine, Harvard Medical School, Boston, Massachusetts, USA; Department of Immunology and Infectious Diseases, Harvard T.H. Chan School of Public Health, Boston, Massachusetts, USA; Department of Pediatrics, Harvard Medical School, Boston, Massachusetts, USA; Division of Global Health, Massachusetts General Hospital for Children, Boston, Massachusetts, USA; Department of Global Health, University of Washington, Seattle, Washington, United States; McGill Genome Centre, McGill University, Montréal, Québec, Canada; McGill Centre for Microbiome Research, McGill University, Montréal, Québec, Canada

## Abstract

Two circular chromosomes are a defining feature of the family *Vibrionaceae*, including the pathogen *Vibrio cholerae*, with rare reports of isolates with a single, fused chromosome. Here we report chromosome fusions in clinical *V. cholerae* O1 isolates, including several independent fusion events stable enough to be transmitted between patients within a household. Fusion occurs in a 12 kilobase-pair homologous sequence shared between the two chromosomes, which may lead to reversible chromosomal fusion.

## Main

Cholera is a waterborne infectious disease affecting millions of people yearly and causing outbreaks where sanitary infrastructure is inadequate^1^. The ongoing seventh cholera pandemic is caused by a pathogenic lineage (7PET) of *Vibrio cholerae* O1 (*Vc*) carrying four virulence-associated genomic islands: VPI-1, VPI-2, VSP-I and VSP-II^2–4^. In Bangladesh, where cholera is endemic, the 7PET sublineage BD2 was dominant between 2009 and 2018, followed by BD1.2, which was responsible for a large outbreak in Dhaka in 2022^5^. *Vc* typically carries two chromosomes: the larger ∼3 megabase-pair (Mbp) chromosome 1 and the smaller ∼1 Mbp chromosome 2. When replication of chromosome 2 is impaired under laboratory conditions, the two chromosomes can fuse to restore cell replication^6^. Out of thousands of sequenced genomes, only three *Vc* with fused chromosomes have been reported to date from natural environments^6–8^. These have typically been considered rare exceptions to the bipartite genome structure. However, due in part to limitations of short-read sequencing, the prevalence of chromosome fusion in *Vc* remains unknown.

Here, we aimed to detect chromosome fusion in clinical *Vc* isolates and identify potential fusion mechanisms. To do so, we used long-read nanopore sequencing of 467 *Vc* isolates, collected between 2015 and 2018 from 47 patients (21 index cases and 26 household contacts) from 21 households in Dhaka, Bangladesh (Fig 1A, Table S1, Fig S1). All isolates were identified as serotype O1. Of these, 400 genomes assembled into two circular chromosomes (3 and 1 Mbp each), 58 into a single 4 Mbp chromosome, and nine were incompletely assembled. All 58 single-chromosome genomes resulted from an apparent fusion of chromosomes 1 and 2. These fused chromosomes were identified in ten different cholera patients from five different households. In three households, fused chromosomes were assembled from both the index cases and household contacts sampled 0-5 days later, suggesting that fused genomes are stable enough to be transmitted (Fig 1B). In the other two households, fusions were detected in only one patient per household.

**Fig. 1.**
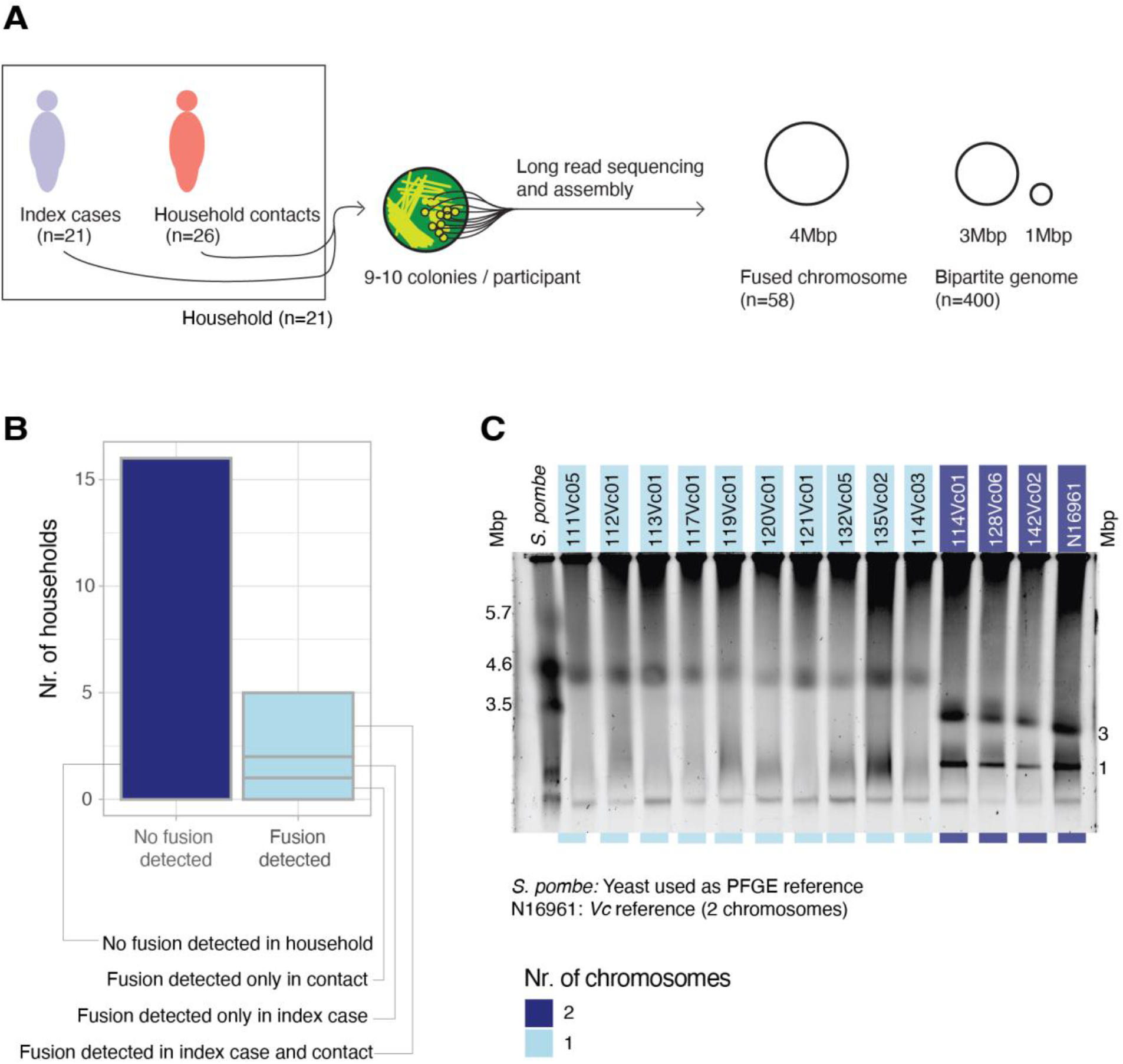
*V. cholerae* with a fused chromosome identified in multiple patients and households. **A:** Schematic representation of the study design; **B:** Occurrence of chromosome fusion by nanopore sequencing in the sampled households; **C:** PFGE results for the putatively fused (light blue) and non-fused (dark blue) chromosomes identified by sequencing. *Schizosaccharomyces pombe* is used as a DNA size marker and *Vc* strain N16961 as a well-characterized isolate with two chromosomes.

As independent verification of chromosome fusion, we subjected a subset of isolates to pulsed-field gel electrophoresis (PFGE). We included one putative fused-chromosome isolate per patient for which at least one fused chromosome was assembled (n=10) and three putative non-fused-chromosome isolates for comparison. As expected, we detected one band at 4 Mbp for all putative fused-chromosome isolates and two bands at 3 and 1 Mbp for the non-fused isolates, corresponding to the known sizes of chromosomes 1 and 2, respectively (Fig 1C). Some of the fused isolates have a weak band at 1 Mbp in addition to the band at 4 Mbp, but the lack of a band at 3 Mbp in these isolates indicates that fusion did occur. Chromosome fusion therefore does not appear to be an artefact of sequencing or assembly.

To understand the mechanism of fusion, we scanned the flanking regions on either side of the fusion site. We found that in fused chromosomes, the chromosome 2 sequence is flanked by a 12 kilobase-pair (Kbp) homologous sequence (HS1) oriented in the same direction on either side of the integrated chromosome 1 sequence. In many non-fused strains, HS1 appears twice: once on chromosome 1 and once on chromosome 2 (Fig 2A). This suggests homologous recombination at HS1 as a potential fusion mechanism. On chromosome 1, HS1 is located within VPI-2 and encodes multiple proteins linked to horizontal gene transfer (Fig 2B). To further support the observation of fusion in the assemblies, we screened the raw sequence data for reads spanning HS1 and its flanking regions, which were highly concordant with the results of the assemblies (Fig S2).

**Fig. 2.**
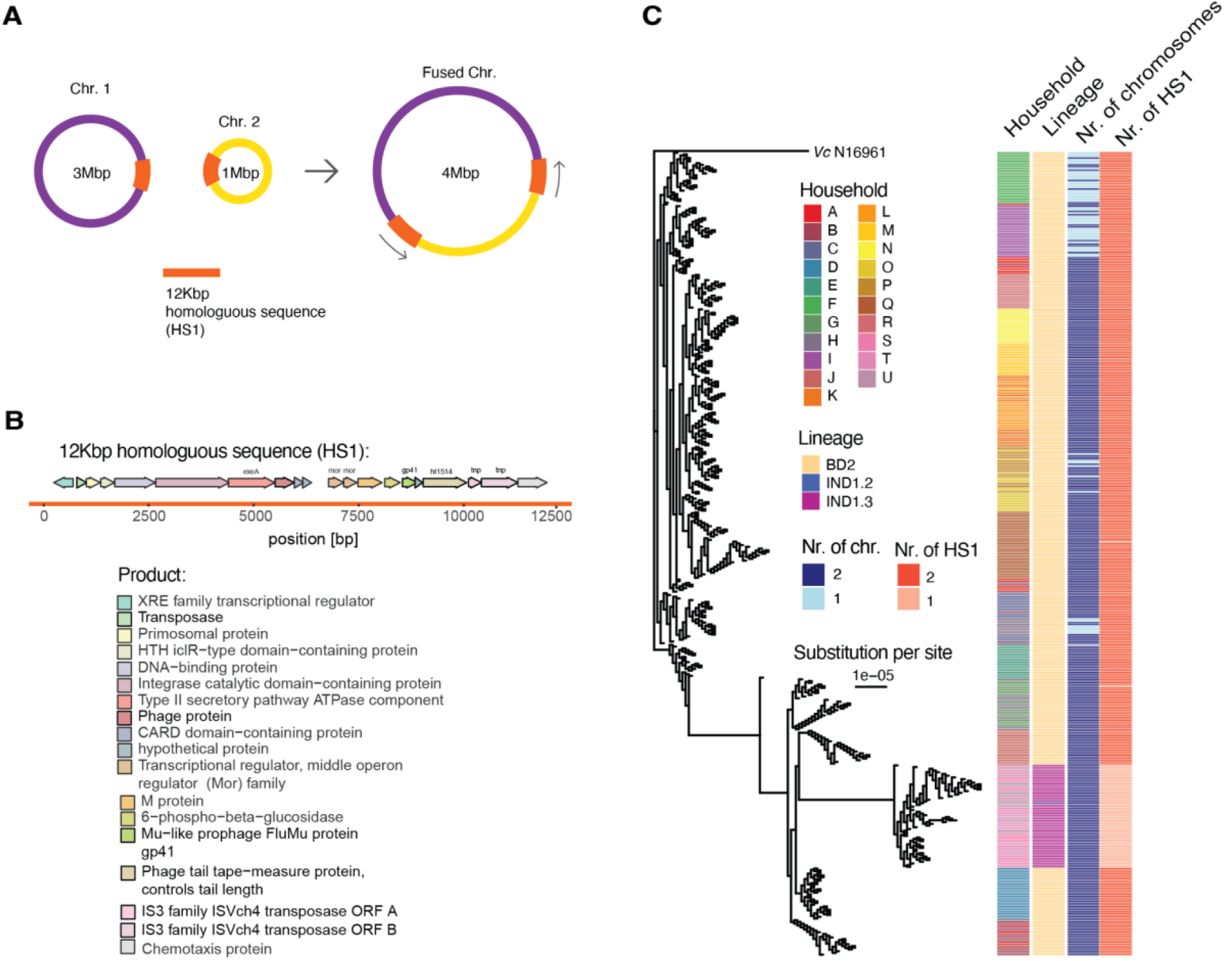
Chromosome fusion events occur in *V. cholerae* sublineages with a shared homologous sequence on each chromosome. **A:** Schematic representation of the chromosome fusion; **B:** Annotation of genes within the homologous sequence (HS1) at the fusion site; **C:** Phylogenetic tree based on 198 high-quality SNVs, along with household membership, the 7PET sublineage designation, the number of chromosomes identified by sequencing, and the number of times HS1 was detected in the genome. Note that branches of length zero are illustrated with a small minimum branch length for clarity.

To investigate the dynamics of chromosome fusion in our samples, we examined the phylogenetic distribution of fused and non-fused states and reconstructed the likely ancestral states along the phylogeny. The phylogenetic clustering of patients within a household is consistent with previous studies^9,10^, suggesting *V. cholerae* transmission within households (Fig 2C), including instances of fused chromosome transmission (Fig 1B). All fused chromosomes were part of the 7PET sublineage BD2 (Fig 2C). All BD2 genomes in our dataset contained two copies of HS1 (one on each chromosome), likely explaining their propensity for fusion. By contrast, other sublineages – notably IND1.3 – contained only one HS1 copy (Fig 2C), preventing chromosome fusion through the same mechanism. Reconstructing the ancestral chromosome states showed that fusion events (n=10) were less common than fission (n=17) (Fig S3A). Independent fusion events were inferred at five nodes in the phylogeny within BD2 (Fig S3B). The closest subsequent fission events were detected at distances of 1-1.9×10^−5^ substitutions/site, corresponding to 41-79 single nucleotide variants (SNVs). Assuming a molecular clock of 3.5 SNVs/year^11^, this suggests that fused chromosomes can remain stable for ∼12-22 years. Yet, we detected closely-related fused and non-fused isolates collected from the same household and patient, suggesting rapid fusion/fission events can occur within patients.

We further investigated the frequency of chromosome fusion more broadly across the order *Vibrionales*, in which a bipartite genome is considered a defining feature. We downloaded publicly available long-read sequences (n=302, 251 of which passed our quality controls), 73.7% (185/251) of which were *Vc* (Fig S4A). Of the fully circular assemblies (n=203), four assembled to one fused chromosome. One of these was a *V. natriegens* genome, whose chromosomes were lab-engineered to be fused^12^. The remaining three were clinical *Vc* isolates^13,14^. Two of these, from the IND1.1 sublineage, contained two directly-oriented copies of HS1, as in our isolates. The third isolate, which was related to IND2, also contained two copies of HS1 but in the opposite orientation, suggesting a local inversion of one HS1 copy. These results suggest that fusion, while rare, can occur in different *Vibrio* species and *Vc* sublineages.

We next asked if HS1 is unique, or if other potential fusion sites exist in the genome. For each circular, non-fused public genome (n=199), we compared chromosome 2 against chromosome 1 for regions of homology. We identified two such regions longer than 10 Kbp in *Vc*, one of which corresponded to HS1 and the other to VSP-I^2^ (Fig S4, Fig S5). Although VSP-I might serve as a potential fusion site, we currently lack evidence for this as none of the fused-chromosome genomes carries more than one VSP-I copy.

Several genes are known to be involved in *Vc* chromosome replication, and these all appear to be present and intact in fused chromosomes. Both *Vc* chromosomes encode two partitioning (*par*) genes involved in separating chromosomes to daughter cells^15^. We detect all four *par* genes in the fused chromosomes sequenced here (n=58), with no mutations compared to *par* genes on non-fused chromosomes. Fused chromosomes also contain origins of replication from both chromosomes (ori1 and ori2) and *crtS* (*C*hr2 *r*eplication *t*riggering *S*ite). *crtS* is located closer to ori1 than ori2 is to ori1 in all fused chromosomes. This arrangement of loci has previously been associated with ori2 being active in a naturally fused *Vc* chromosomes^16^. Finally, *Vc* can be engineered to support fusion of chromosomes through deletion of the DNA adenine methylase Dam^6^. The *dam* gene was present in all our genomes with no non-synonymous mutations detected; therefore loss of *dam* function is unlikely to explain the observed fusions. Further experiments will be needed to understand how fused chromosomes replicate, and whether fusion affects bacterial growth or other phenotypes.

Together, our results show that chromosome fusion via homologous recombination is more prevalent and potentially more stable than previously thought. The clinical or phenotypic consequences of fusion, if any, remain to be explored. This study reveals chromosome fusion in clinical *V. cholerae* O1 strains at an unprecedented scale and highlights the power of long-read sequencing to identify structural variation in bacterial genomes.

