## Supplementary material for "Prevalent chromosome fusion in *Vibrio cholerae* O1": Methods

### Ethical statement

The Ethical and Research Review committees of the icddr,b (approval number PR-11041) and the Institutional Review Boards of Massachusetts General Hospital, the University of Washington (deemed not human study due to all samples de-identified) and McGill University (A07-M43-21B (21-07-026)) approved the study. All adult subjects in the study provided written informed consent and the parents/guardians of children provided written informed consent.

### *V. cholerae* isolate collection

Stool and rectal swab were sampled from patients with cholera admitted to the International Centre for Diarrhoeal Disease Research, Bangladesh (icddr,b) Dhaka Hospital and from their household contacts, as described in prior studies[1]. Patients presenting to the hospital with severe acute diarrhoea and a stool culture positive for *V. cholerae* O1 were considered index patients. Persons who shared the same cooking pot with an index patient for 3 or more days are considered household contacts and were enrolled within 6 hours of the presentation of the index patient to the hospital. Rectal swabs were collected daily from household contacts during a 10-day period after presentation of the index case. Household contacts underwent daily clinical assessment of symptoms. Household contacts were defined as infected if any rectal swab culture was positive for *V. cholerae* O1. *V. cholerae* serotypes were determined using slide agglutination testing with polyvalent and specific antisera as in prior studies[2].

We excluded patients below 2 years of age and above 60 years old or with major comorbid conditions[3,4]. Rectal swabs and stool from the day of enrollment and follow-up time points were collected and placed immediately on ice after collection and stored at -80°C until DNA extraction.

*V. cholerae* isolates were cultured from stool samples from culture-positive participants. Stool samples were streaked directly onto tellurite taurocholate gelatin agar, a selective medium for *V. cholerae*, and incubated at 37°C for 18-24 hours. Ten colonies from each participant were selected and inoculated into Luria-Bertani (LB) broth and grown at 37°C overnight. Liquid cultures were used to make 30% glycerol stocks and stored at -80°C. Frozen glycerol stocks were then shipped to the University of Washington.

### *V. cholerae* DNA extraction

*V. cholerae* from frozen glycerol stocks were streaked onto LB agar plates and incubated at 30°C for 24 hours. One colony per plate was used for liquid culturing in LB broth and incubated at 30°C for 18 hours with agitation. DNA was extracted from saturated liquid cultures using the DNeasy Blood and Tissue 96-kit (Qiagen) according to the manufacturer's protocol. Briefly, *V. cholerae* samples were pelleted and resuspended in Buffer ATL and treated with proteinase K at 56°C for 30 minutes followed by RNase A (Qiagen) at room temperature for 5 minutes and Buffer AL for an additional incubation at 56°C for 10 minutes. Samples were then treated with 100% EtOH and transferred to Qiagen DNA spin columns. DNA was washed using Buffer AW1 and Buffer AW2. Purified DNA was eluted in 100 µL 10 mM Tris-HCl and stored at -80°C until ready for sequencing.

### DNA sequencing

The extracted DNA was prepared for sequencing using the Nanopore Rapid Barcoding 96 v14 (Oxford Nanopore Technologies) with approximately 200ng of purified DNA per sample to generate sequencing libraries. The libraries were sequenced on a R10.4.1M PromethION flow cell. Raw sequencing data was basecalled and demultiplexed to FASTQ files using the Dorado basecaller integrated into MinKNOW v 23.07.5 (Oxford Nanopore Technologies) using the model dna_r10.4.1_e8.2_400bps_sup@v4.2.0 with read splitting, adapter trimming and barcode trimming enabled in the basecaller. Read data generated for this study is available on NCBI SRA (BioProject PRJNA1121190).

### Sequence analysis of isolates collected for this study

#### Assembly, Quality Control and Annotation

Reads of each sample were assembled using Flye[5] in –nano-hq and deterministic mode. We used dnaapler[6] to reorient the contigs such that all chromosomes start with the initiating codon of *dnaA* if present (chromosomes 1 and fused chromosomes) or with *repA* homologous sequences if *dnaA* was not present (chromosomes 2).

We included the reference strain *Vc* N16961 as a control, which assembled in two chromosomes, as expected. To further reduce the chances of misassembly, we manually assembled the sequences of 13 isolates using Trycycler[7]. This subset included one putative fused-chromosome isolate per patient for which at least one fused-chromosome was assembled (n=10) and three non-fused chromosome isolates as references. We followed the Trycycler workflow. First, the reads of each sample were filtered using Filtlong[8] (minimum length 1000bp and excluding the worst 5% of read bases) and subsampled to 12 files. Of each isolate, 3 subsets were assembled with Flye[5], Miniasm[9] and Minipolish[10], Raven[10,11] and Canu[10–12]. All assembled contigs were clustered per sample, aligned and manually curated. We considered contig clusters as valid, if they (i) occurred in at least two assemblies and were of similar size, (ii) no kilobase of sequence was below the identity threshold of 25% when compared to the rest of the contig and (iii) could be circularly assembled. Of each valid contig cluster, a consensus was built which together form the assembly for each sample. The assemblies resulting from the Trycyler workflow confirmed the chromosome fusion state observed from the Flye assemblies. We used the Flye assemblies for all isolates for all subsequent analysis. We used Metaphlan2[13] to screen for contaminations in our samples and Nanostat[14] to assess the quality of our assemblies. All assemblies were annotated using Bakta[13,15].

#### Phylogeny

We called variants of each of our isolates to the *V. cholerae* reference genome N16961 (GCF_001250235.2) using Medaka[16] with a minimal quality score threshold of 40 and excluding indels using vcftools[17]. We identified varying sites from the resulting consensus sequences using snp-sites[18] and constructed a maximum-likelihood phylogenetic tree using RAxML[19] (v8.2.12, model GTRCAT).

#### Sub-lineage assignment

We aimed to assign our isolates to one of the sub-lineages within the clonal lineage causing the current 7th pandemic (7PET). We selected one previously assigned and publicly available genome per sub-lineage[20] as reference and called variants of each of our sequences to each reference using the Medaka variant caller[16]. Each isolate was assigned to the sub-lineage to whose reference the smallest number of high quality variants was identified (using a minimal quality score threshold of 40).

**Identification of HS1**

To identify potential fusion sites, we used blastn[21], identifying sequences, which are shared between chromosome 1 and chromosome 2. We detected a 12 Kbp long sequence which was identical between chromosome 1 and chromosome 2 in most strains (HS1). We screened for the occurrence of HS1 in all our genomes using blastn[21].

To further substantiate the observed chromosome fusion, we next aimed to identify reads which span HS1 and are either flanked by chromosome 1 or chromosome 2 sequences (non-fused case) or are flanked by sequences of each chromosome on each side (fused case). To do so, we mapped the reads of each sample to a one-chromosome assembly (135Vc06) and a two-chromosome assembly (135Vc05). From the bam file, we extracted reads which span the HS1 and additional 500 additional bp on each side using samtools. We excluded secondary mappings and mappings with large insertions (>1000 bp).

We excluded reads in which a sequencing adapter sequence was identified, as these could potentially be artifactual hybrid molecules. To do so, we converted the reads to fasta files and screened for adapter sequences using blastn. All reads in which an adapter sequence with > 95% coverage and identity was identified were excluded from further analysis.

**Ancestral state reconstruction**

To infer the ancestral chromosome state (non-fused vs. fused) of our isolates, we used the R package ape[22] and transition finder of the package Quidiphydi[23] to identify fusion and fission events.

**Screening of genomic islands**

We queried our genomes for the presence and location of the virulence associated genomic island VSP-I by screening for the known flanking regions[24] using blastn[21]. We extracted the sequence between the flanking sides using bedtools getfasta[24,25].

**Comparison of sequences involved in chromosome replication**

*Vc* typically encodes two sets of partitioning (*par*) genes (*parAB* on chr1 and *parAB2* on chr2), which are involved in separating the chromosome molecules to daughter cells. As chromosome 2 can be lost upon depletion of *parAB2*, we hypothesised that *parAB2* might be lost or mutated in strains with a fused chromosome. To test this, we downloaded the amino acid sequence of reference *Vc par* proteins (QEO41389.1, QEO42567.1, AAF97006.1, and AAF97005.1 for ParA, ParB, ParA2 and ParB2, respectively) and screened their occurrence in our genomes using tblastn[21].

To compare the *dam* gene sequence and the presence of the origins of replication on the fused chromosomes, we examined the bakta annotation. We screened for *crtS* (*C*hr2 *r*eplication *t*riggering *S*ite)[26] in our genomes using blastn[21].

### *V. cholerae* plug preparation and pulsed-field gel electrophoresis (PFGE)

*V. cholerae* isolates from frozen glycerol stocks were cultured as described above. *V. cholerae* plugs were created as previously described (CDC Pulsenet protocol)[27]. Briefly, *V. cholerae* cells were inoculated into a cell suspension buffer (100 mM Tris, 100 mM EDTA, pH 8.0) to an optical density of 2.0. Plugs were caste using Proteinase K (ThermoFisher) and 1% SeaKem Gold agarose (Lonza) prepared in Tris-EDTA (TE) buffer (10 mM Tris, 1 mM EDTA, pH 8.0). Plugs were lysed in cell lysis buffer (50 mM Tris, 50 mM EDTA, pH 8.0 with 0.1% Sarcosine (Sigma) and 0.1 mg/mL Proteinase K (ThermoFisher)), incubated at 55°C for 30 minutes with shaking at 200 rpm. Plugs were then washed twice with sterile, distilled water and four times with TE buffer. Plugs were stored in TE buffer at 4°C until PFGE was performed.

Gels, 21cm long, were caste for PFGE using 0.8% SeaKem Agarose in 1X Tris acetate EDTA (TAE) buffer. PFGE was performed on a CHEF-DR III system (Biorad) with runtime of 60 hours, at 2 Volts/cm with 30 min switch time (initial and final) at a 106° angle with 1X TAE running buffer chilled to 14°C. The gel was then stained using SYBR Gold (Invitrogen) and visualised using Amersham Typhoon 5 (Cytiva). *Schizosaccharomyces pombe*[28] DNA was used as a DNA marker.

### Sequence analysis of publicly available sequences

#### Selection of publicly available sequences

To contextualise the sequences acquired for this study, we used two different datasets, which we compiled from publicly available sequences: (a) long-read sequenced genomes of the order *Vibrionales* and (b) short read sequenced genomes of the species *V. cholerae*.
Dataset (a) consisted of all *Virbrionales* genomic sequences that were acquired using Oxford Nanopore Technology (ONT) or Pacbio and available from NCBI SRA on the 19th of December 2023 (n=302).
We compiled dataset (b) aiming to reflect the diversity within clinical *V. cholerae* and to focus our study period (2015-2018) and geographic region of isolation (Bangladesh) for comparison. To do so, we downloaded the following sets of short reads:
(b.i) Clinical *V. cholerae* overview: from NCBI pathogens, we selected *V. cholerae* which were collected between the 1st of January 2000 and the 18th of January 2024 and included a maximum of three samples per year and country (n=790, 457 of which could be successfully downloaded using NCBI Batch entrez).
(b.ii) Diversity within *V. cholerae* O1: We downloaded a previously compiled selection of *V. cholerae* O1[29] (all O1 isolates from this study).
(b.iii) 7PET diversity from South and East Asia: Of a previously compiled 7PET set of genomes[30], we included strains which were isolated between 2000 and 2018 in Eastern Asia, Southeast Asia and South Asia. Moreover, we included the sequences acquired in this[30] and another study conducted in Bangladesh around our study period [31].

#### Assembly, Quality Control and Annotation

Long-read sequences were assembled using Flye[5] (--nano-raw mode) and the species was identified using GTDB-tk[32]. We controlled the quality of the assemblies using Kraken2[33] / Bracken[34], as well as Nanostat[14]. We excluded sequences which were not identified as part of the order *Vibrionales*, where >10% of the contigs were not assigned as the most abundant genus (suggesting contamination), whose total assembly length was smaller than 3.5 or larger than 7 MB, or which had an average read depth below 5X, resulting in 251 genomes of 24 different species.
Short-reads were trimmed using Trimmomatic[35] and were assembled using Spades[36] via Unicycler[37]. We assessed the quality of the resulting assemblies using Quast[38] and used

Metaphlan2[13] to screen for possible contaminations. We excluded sequences which had a Metaphlan purity below 95%, an average read depth below 20X, an average read quality below 75, where the resulting assembly was shorter than 3.65 or longer than 4.5 MB or had N50 value below 2000. This resulted in a set of high quality *V. cholerae* short-read genomes (n=1,223). The accession numbers of all publicly available sequences analysed in this study can be found in Table S2.

**Phylogeny**

For the *Vibrionales* long reads, we built a SNP-based phylogeny, as described above. Briefly, we called variants of each sample to the *V. cholerae* reference genome N16961 (GCF_001250235.2) using Medaka[16] (minimal quality threshold 40), identified varying sites of the resulting consensus sequence using snp-sites[18] and built a phylogenetic tree using RaXML [19] (v8.2.12, model GTRCAT).

We built a core genome for the *V. cholerae* short reads using panaroo[39] and aligned it using mafft[40]. Variant sites were identified using snp-sites[18] and a phylogenetic tree was constructed using fasttree[41].

**(Sub)-Lineage Assignments**

We used 'Is it 7PET' [42] to identify which publicly available *V. cholerae* sequences belonged to the 7PET lineage. To identify the sub-lineages within the 7PET, we quantified the number of SNV to one reference per sub-lineage each, and assigned the sub-lineage with the least number of SNV, similar as described above. SNV were called using Freebayes[43] via snippy[44] (minimum read depth = 10, minimum fraction = 0.7 and minimum quality = 100).

**Copy Number Estimation of Potential Fusion Sites**

We used blastn[21] to identify the presence and location of the potential fusion sites HS1 and VSP-I in the publicly available *V. cholerae* genomes (>95% identity and >90% coverage). We used bwa[45] to align the reads of each isolate to its respective assembly. From the resulting bam files, we retrieved the average read depth of the genome and from the specific potential fusion sites using samtools depth[46].

### Data and Code availability

Read data generated for this study is available on NCBI SRA (BioProject PRJNA1121190). Software code used to analyse the data presented in this study is available on GitHUb (https://github.com/acuenod111/Single_chromosome_Vc) and the files to reproduce the analysis and figures can be accessed via the OpenScienceFoundation (https://osf.io/xyfvg/).


### Acknowledgements

We are grateful to the people of Dhaka, where our study was undertaken, to the field, laboratory, and data management staff, who provided a tremendous effort to make the study successful, and to the people who provided valuable support in our study.

### Funding

AC was supported by a Postdoc.Mobility Fellowship from the Swiss National Science Foundation (P500PB_214356). BJS and PL were supported by a Canadian Institutes for Health Research (CIHR) Project Grant and Fellowship, respectively. This work was also supported by a grant from the U.S. National Institutes of Health/NIAID AI106878 (ETR, FQ), K08AI123494 (AAW), and T32HD007233 (DC). RWH and GRS were supported by research grant R35 GM118120 from the National Institutes of Health of the United States of America.

### Author contributions

AIK, FC, SBC, ETR, JBH, RCL, TRB and FQ conducted the study in Dhaka, Bangladesh and collected the patient samples.
DC extracted the DNA for all samples.
PL developed the sequencing workflow, sequenced all samples and compiled the initial assemblies.
AC conducted all other bioinformatic analysis and generated the figures.
DC, RWH, SMM and AAW performed the PFGE assay.
AAW, BJS and PL supervised the project
ETR, FQ, DC, GRS, BJS, AAW, PL, and AC acquired funding for this project.
AC and BJS wrote the original manuscript.
DC, AIK, FC, RWH, SMM, SBC, ETR, JBH, RCL, TRB, GRS, FQ, PL and AAW reviewed and approved the manuscript.

in DNA sequences from a metagenomics sample. *GitHub* https://github.com/jenniferlu717/Bracken.

35. Bolger, A. M., Lohse, M. & Usadel, B. Trimmomatic: a flexible trimmer for Illumina sequence data. *Bioinformatics* **30**, 2114–2120 (2014).
36. Bankevich, A. *et al.* SPAdes: a new genome assembly algorithm and its applications to single-cell sequencing. *J. Comput. Biol.* **19**, 455–477 (2012).
37. Wick, R. R., Judd, L. M., Gorrie, C. L. & Holt, K. E. Unicycler: Resolving bacterial genome assemblies from short and long sequencing reads. *PLoS Comput. Biol.* **13**, e1005595 (2017).
38. Gurevich, A., Saveliev, V., Vyahhi, N. & Tesler, G. QUAST: quality assessment tool for genome assemblies. *Bioinformatics* **29**, 1072–1075 (2013).
39. Tonkin-Hill, G. *et al.* Producing polished prokaryotic pangenomes with the Panaroo pipeline. *Genome Biol.* **21**, 180 (2020).
40. Katoh, K. & Standley, D. M. MAFFT multiple sequence alignment software version 7: improvements in performance and usability. *Mol. Biol. Evol.* **30**, 772–780 (2013).
41. Price, M. N., Dehal, P. S. & Arkin, A. P. FastTree 2--approximately maximum-likelihood trees for large alignments. *PLoS One* **5**, e9490 (2010).
42. GitHub - amberjoybarton/is-it-7pet: Identifying whether a cholera sample belongs to the seventh pandemic El tor (7PET) sub-lineage based on sequencing data or an assembly. *GitHub* https://github.com/amberjoybarton/is-it-7pet.
43. Garrison, E. & Marth, G. Haplotype-based variant detection from short-read sequencing. (2012).
44. GitHub - tseemann/snippy: :scissors: Rapid haploid variant calling and core genome alignment. *GitHub* https://github.com/tseemann/snippy.
45. Li, H. Aligning sequence reads, clone sequences and assembly contigs with BWA-MEM. (2013).
46. Danecek, P. *et al.* Twelve years of SAMtools and BCFtools. *Gigascience* **10**, giab008 (2021).
