## Supplementary Figures for "Prevalent chromosome fusion in *Vibrio cholerae* O1"

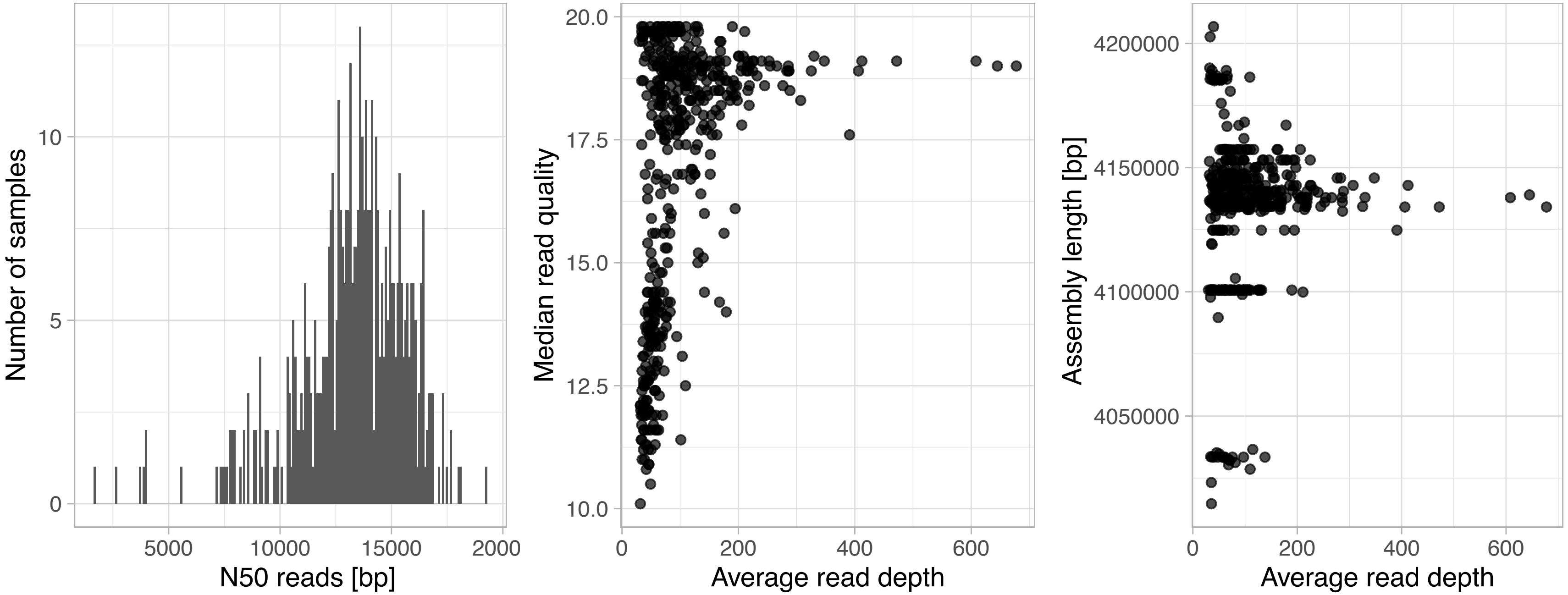


**Figure S1:** Quality measures of the genomes sequenced for this study. **A:** N50 of the reads per sample; **B:** Median read quality and average read depth; **C:** Assembly length and average read depth.

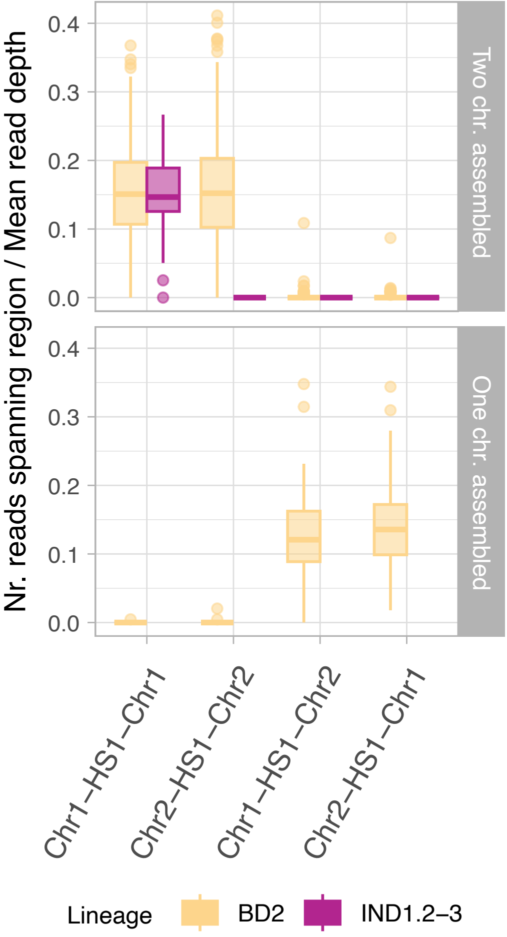


**Figure S2:** Number of reads spanning HS1 and 500bp flanking sequences on either chromosome (Chr1 or Chr2). The chromosomal structure inferred using the assembly is highly concordant with the reads, with reads spanning 'Chr1-HS1-Chr2' or 'Chr2-HS1-Chr1' being found almost exclusively in one-chromosome (fused) assemblies. An exception was the isolate 112Vc05, which assembled into two chromosomes, but for which we identified reads anchored on both chromosomes as well as one read anchored in chromosome 1 on both sides (i.e. a false-negative fusion event in the assembly). All other nine isolates from the same patient assembled into one fused chromosome. The outlier, 112Vc05, likely also contains a fused chromosome, although we cannot exclude a mixed population of fused and non-fused genomes in this colony.

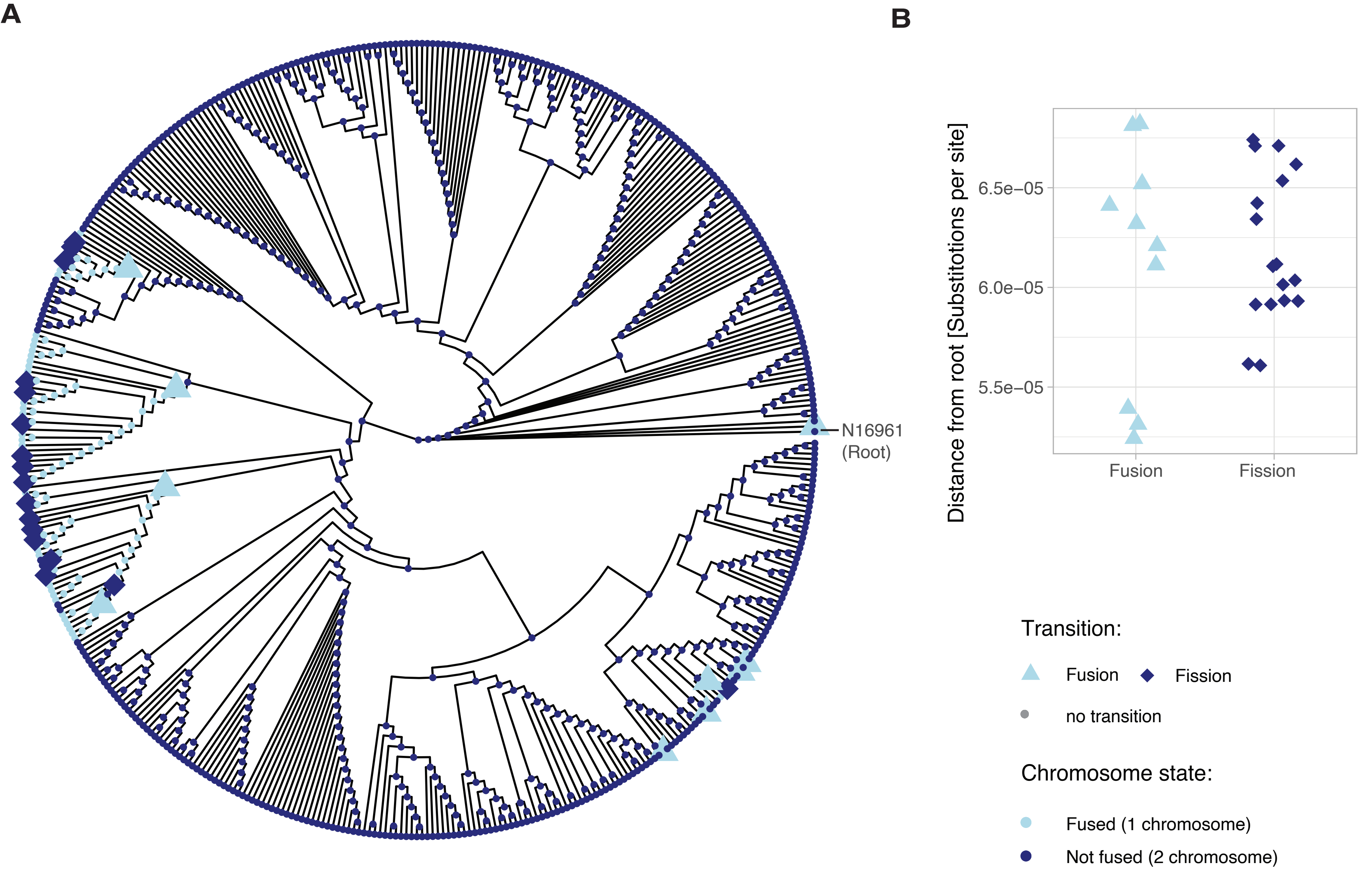


**Figure S3:** Ancestral state reconstruction of the chromosome state (fused or non-fused) in the genomes sequenced for this study (n=467); **A:** Rooted phylogenetic tree (branch lengths not to scale) indicating the reconstructed state for each node. Note that the state of the root (*V. cholerae* N16961) is non-fused, but a root-adjacent deep branch is fused. **B:** Distance from the root to each node where a transition (fusion or fission) is inferred.

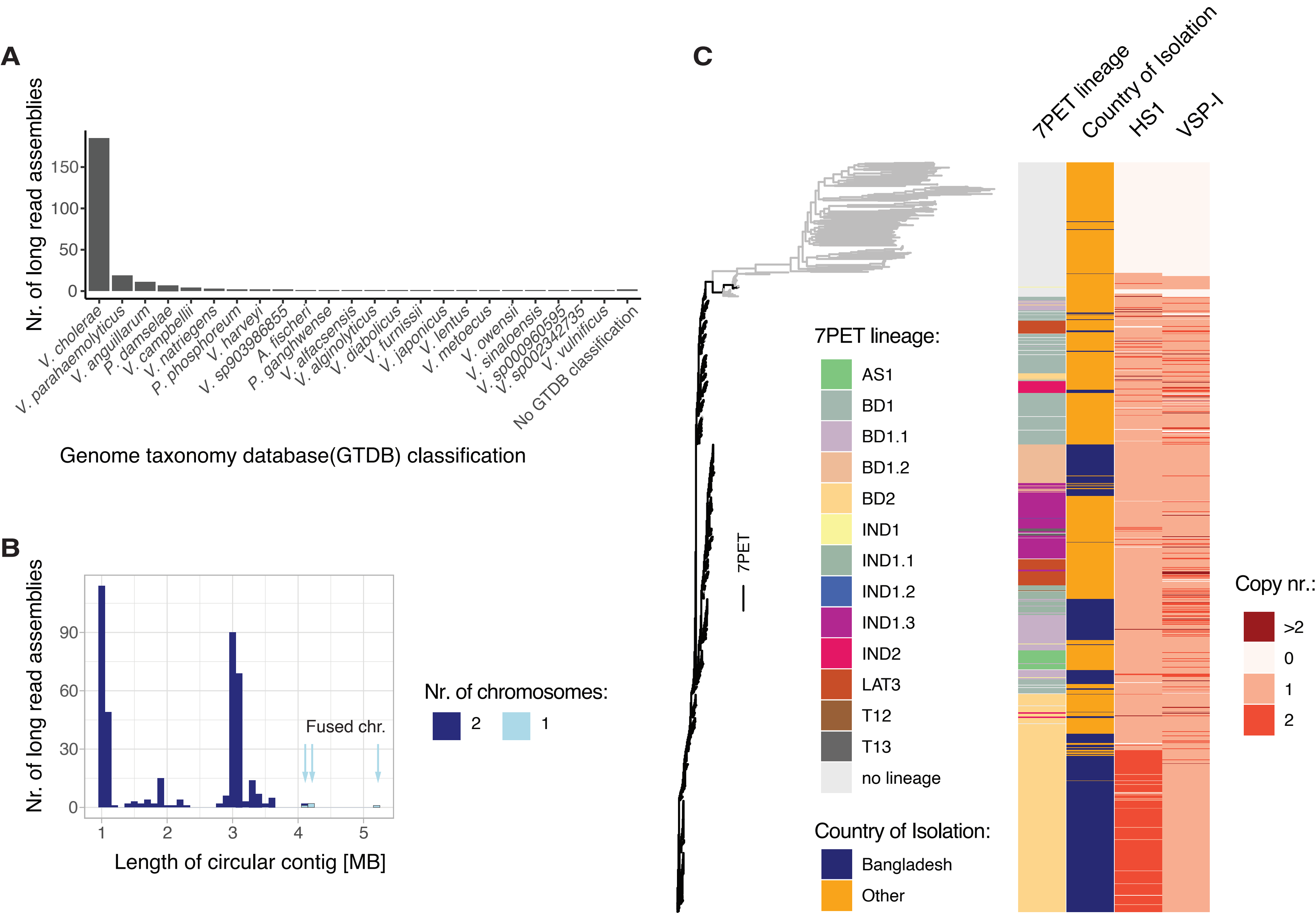


**Figure S4:** Chromosome fusion is rarely detected in other *Vibrio* genome sequences. **A:** *Vibrionales spp.* for which publicly available long-read assemblies were analysed **B:** Contig (chromosome) lengths of fully circularised genomes. Light blue arrows indicate fused chromosomes **C:** Core genome phylogeny of 1,223 publicly available short-read sequenced *Vc* genomes with heatmap indicating which 7PET lineage was assigned and the country of isolation and copy number estimations for HS1 and VSP-I. Whereas HS1 appears predominantly duplicated in the BD2 sublineage, VSP-I is more frequently duplicated in other 7PET sublineages.

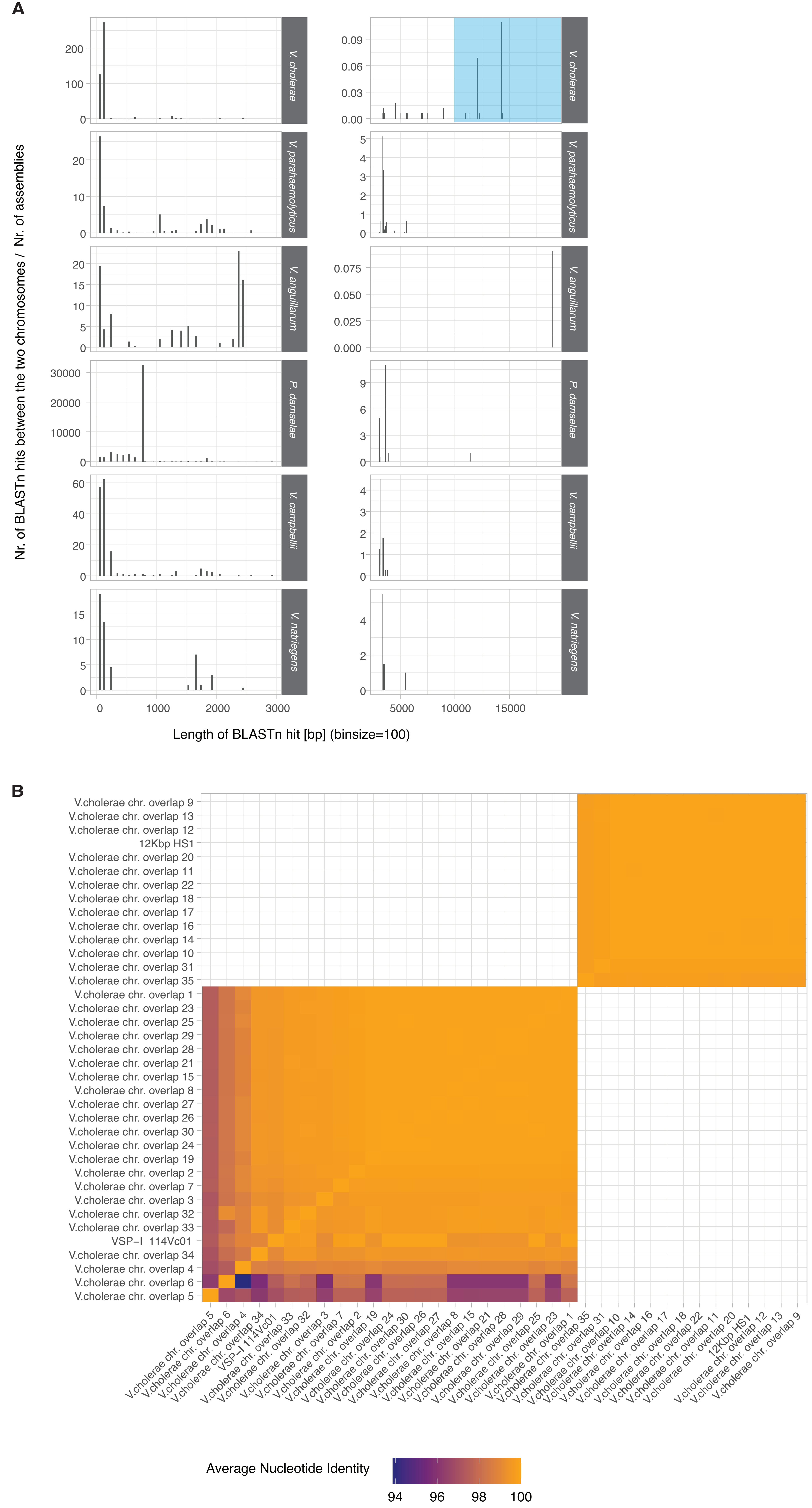


**Figure S5:** Chromosome comparison in bipartite genomes **A:** Number and length of BLAST hits with > 90% nucleotide identity between the chromosomes of each species. Left column: BLAST hits shorter than 3000 bp; right column: BLAST hits larger than 3000 bp. BLAST hits can be overlapping. Highlighted in blue are BLAST hits for *V. cholerae* longer than 10,000bp, which are included in B; **B:** Average Nucleotide Identities of the *V. cholerae* blast hits longer than 10,000bp and HS1 and VSP-I as references.
